# Molecular and morphological analyses of the genus Episoriculus with the description of a new subgenus

**DOI:** 10.1101/2022.09.19.508619

**Authors:** Yingxun Liu, Xuming Wang, Kai He, Tao Wan, Rui Liao, Chen Shunde, Shaoying Liu, Bisong Yue

## Abstract

Shrews in the genus *Episoriculus* are among the least known mammals in China. It occurs mainly in the Himalayas and the Hengduan Mountains. We report the sequencing and analyses of one mitochondrial gene (Cyt *b*) and three nuclear genes (Apob, Brca1, and Rag2) for 77 specimens, and analyses of morphological measurements for 56 specimens. Phylogenetic and morphological studies answer three longstanding questions. First, *Episoriculus sacratus* and *E. umbrinus* are valid species, and not subspecies of *E. caudatus*. Second, *Pseudosoriculus fumidus* is a valid taxon that does not belong to the genus *Episoriculus*. Third, the genus *Episoriculus* has eight valid species: *E. baileyi, E. caudatus, E. leucops, E. macrurus, E. sacratus, E. soluensis, Episoriculus* sp., and *E. umbrinus*. Ultimately, we erected subgenus *Longacauda* subgen. nov. for morphologically distinct *E. macrurus*, which leaves the nominate subgenus *Episoriculus* with the remaining seven congeners.

## Introduction

The genus *Episoriculus* was originally established as a subgenus of *Soriculus* distributed in Southwest China, India, Nepal, and Vietnam [1–3]. Some scholars also believe that *Episoriculus* is a subgenus of *Soriculus* [4–9]. While Repenning [10] elevated *Episoriculus* to a genus due to significant differences in teeth from *Chodsigoa* and *Soriculus* and Jameson and Jones [11] followed this arrangement. And Hutterer [12] considered there are significant differences in morphology between *Episoriculus* and *Soriculus*, which should be an independent taxon. Hutterer [13] promoted recognition of *Episoriculus* as an independent genus.

The taxonomic status of genus *Episoriculus* has changed through time (Table 1). Allen [14] described *S. macrurus, S. caudatus sacratus*, and *S. caudatus umbrinus*. Ellerman and Morrison-Scott [1] erected *Episoriculus* as a subgenus of *Soriculus*, which included *S. leucops* and *S. caudatus* (including five subspecies: *S. c. baileyi, S. c. caudatus, S. c. fumidus, S. c. sacratus*, and *S. c. umbrinus*). Honacki *et al*. [6] considered *Episoriculus* containing *S*. (*E*.) *leucops*, *S*. (*E*.) *macrurus*, *S*. (*E*.) *caudatus*, and *S*. (*E*.) *fumidus*, and elevated *S. caudatus. baileyi* and *S. caudatus. fumidus* to species. Hoffmann [7] also recognized that *Episoriculus* contained four species, but it was not completely consistent with Honacki *et al*. [6]: *S*. (*E*.) *leucops*, *S*. (*E*.) *macrurus*, *S*. (*E*.) *caudatus*, and *S*. (*E*.) *fumidus*, and relegated *S. baileyi* to a subspecies of *S*. (*E*.) *leucops*. Corbet and Hill [8] and Hutterer [9] followed this arrangement. Hutterer [13] listed four species being consistent with Hoffmann [7]. Motokawa and Lin [13] elevated *S. baileyi* to a species based on morphology. Based on the karyotypes and overall differences in skull, Motokawa *et al*. [14] stated that *E. caudatus* appeared to include two species: the larger *E. caudatus* and smaller *E. sacratus* (with the subspecies: *E. s. soluensis* in Nepal and Sikkim, *E. s. umbrinus* in Assam, Myanmar, and Yunnan, China, and *E. s. sacratus* in Sichuan, China). He *et al*. [15] noted that *P. fumidus* did not belong to *Episoriculus*. Abramov *et al*. [2] considered *Episoriculus* to be comprised of *E. baileyi, E. caudatus, E. leucops, E. macrurus*, *E. sacratus*, *E. soluensis*, and *E. umbrinus*, while *E. fumidus* was transferred to new genus *Pseudosoriculus* as *P. fumidus*. Mittermeier and Wilson [3] considered *Episoriculus* to include eight species, including *P. fumidus*. Thus, the taxonomic status of *E. caudatus*, *E. leucops*, and *E. macrurus* has been stable, but the recognition of *P. fumidus, E. sacratus*, *E. umbrinus*, *E. baileyi*, and *E. soluensis* remains controversial.

**Table 1.** Major classification systems of the genus *Episoriculus*. Genus name (subgenus name), recognized species (subspecies) are presented, but not synonyms.

| Allen | Ellerman &<br>Morrison-Scott | Hoffmann | Hutterer | Mittermeier<br>& Wison |
| --- | --- | --- | --- | --- |
| *S. caudatus* | *S. caudatus* | *S. caudatus* | *E. caudatus* | *E. caudatus* |
| *(umbrinus)* | *(umbrinus)* | *S. leucops* | *E. macrurus* | *E. umbrinus* |
| *S. leucops* | *(baileyi)* | *(baileyi)* | *E. leucops* | *E. sacratus* |
|  | *(sacratus)* | *S. macrurus* | *E. fumidus* | *E. leucops* |
|  | *(fumidus)* | *S. fumidus* |  | *E. macrurus* |
|  | *S. leucops* |  |  | *E. soluensis* |
|  |  |  |  | *E. fumidus* |
|  |  |  |  | *E. baileyi* |

To clarify the taxonomic status of the species assigned to *Episoriculus*, we assembled a collection of *Episoriculus* from different regions of China, from 2001 to 2020. In the study, 77 specimens were used for morphometric analyses, and 56 specimens used for geometric morphometric analyses. With reference to the methods described by He *et al*. [15], Chen *et al*. [18–19], we amplified a mitochondrial gene (Cyt *b*) and three nuclear gene fragments (Apob, Brca1, Rag2). Herein, we explore the phylogenetic relationships and taxonomic status of these species based on molecular and geometric morphometric data.

## Materials and methods

### Ethics statement

All samples were obtained following the regulations of China for the implementation of the protection of terrestrial wild animals (State Council Decree [1992] No. 13). Collecting was approved by the Ethics Committee of Sichuan Academy of Forestry (no specific permit number). Voucher specimens were deposited in Sichuan Academy of Forestry, Chengdu, China. Sampling and sequencing A total of 77 specimens were collected in China, including 31 individuals of *E. macrurus*, 20 individuals of *E. caudatus*, 4 individuals of *E. sacratus*, 5 individuals of *E. umbrinus*, 4 individuals of *E. soluensis*, and 10 individuals of *E. leucops;* no samples of *E. baileyi* and *P. fumidus* were available (Table 2, Fig. 1). All specimens were identified based on external characteristics according to Smith and Xie [20]. Voucher specimens were deposited in the Sichuan Academy of Forestry. We also collected muscle and liver tissue in 95% ethanol, which were subsequently stored at −75 °C for molecular studies.

**Fig. 1.**
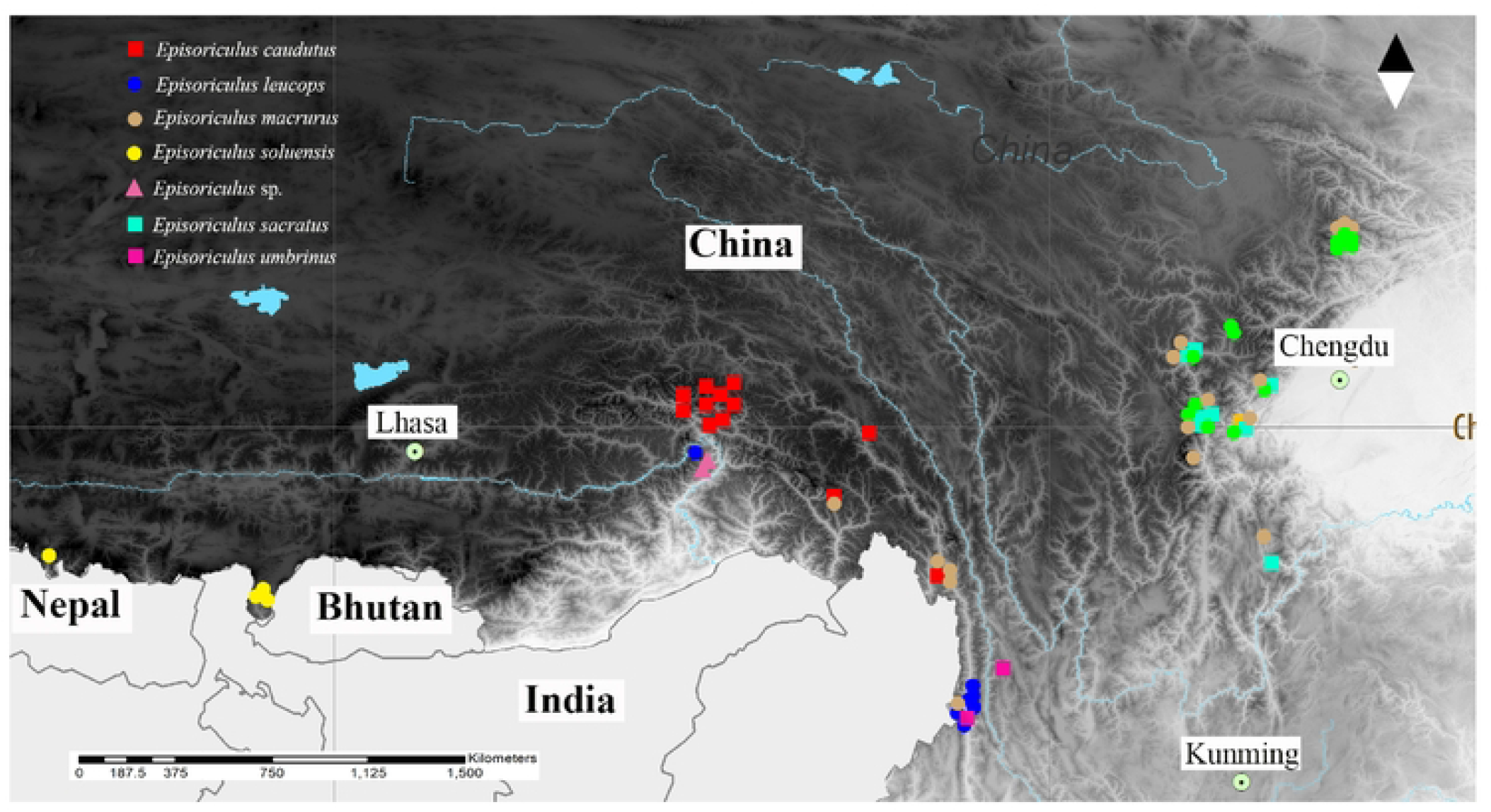
Map of genus *Episoriculus*, showing localities sampled for this study. This is the Fig 2 legend.

**Table 2.**
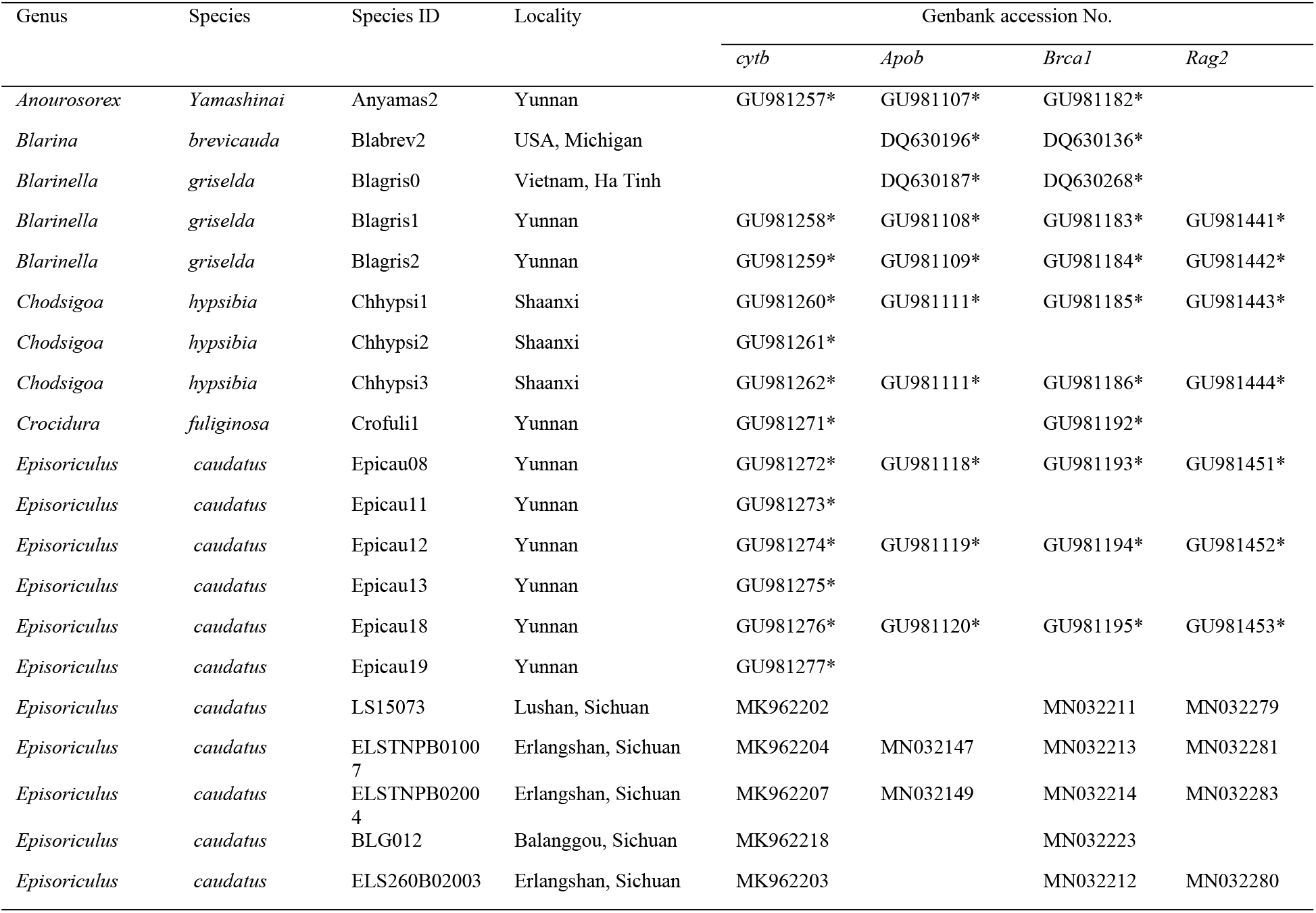
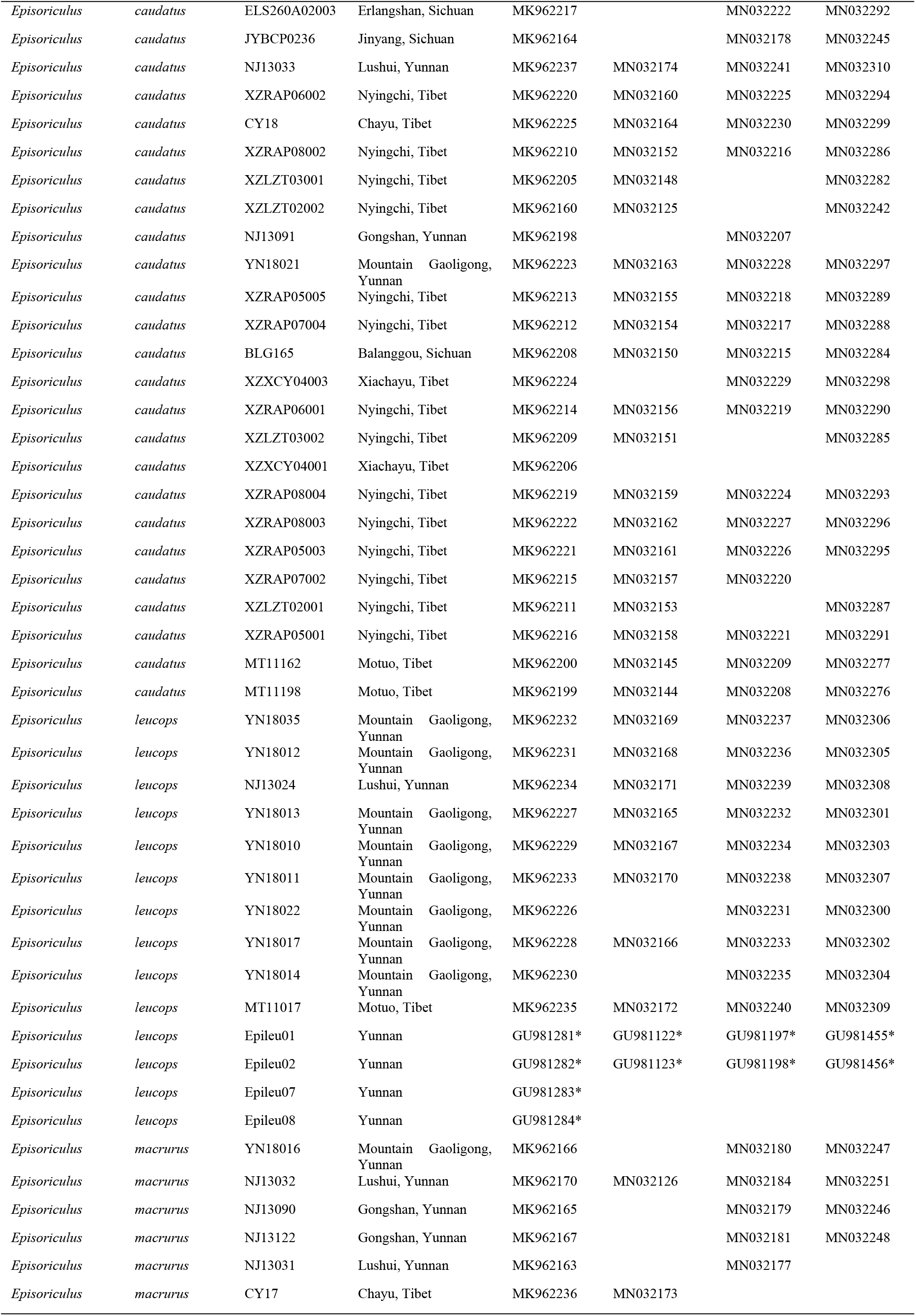
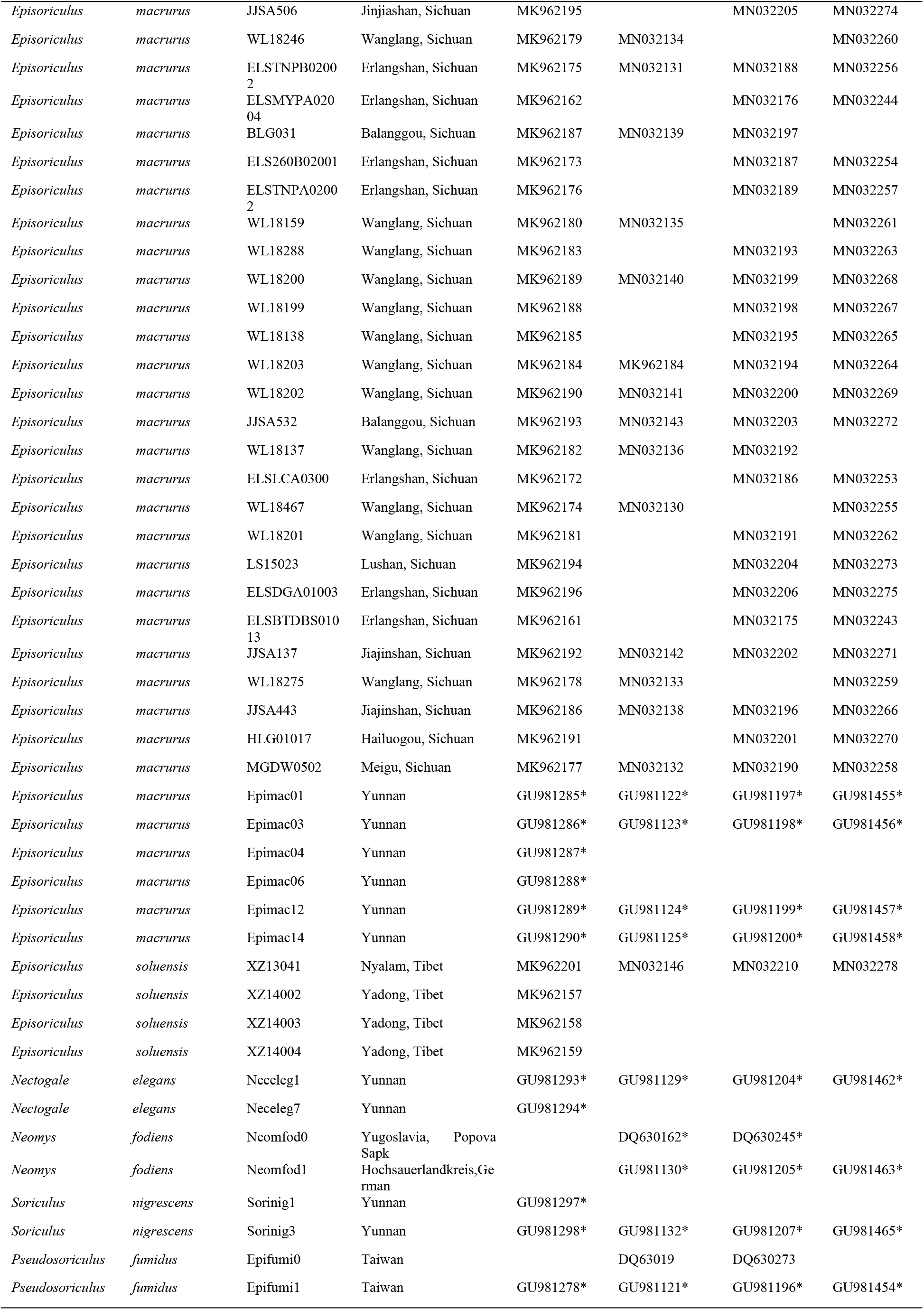
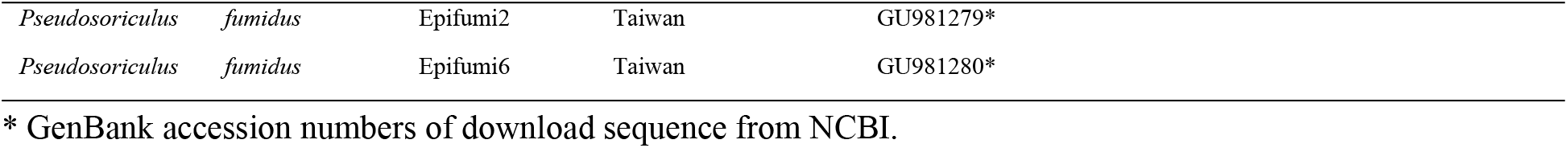
Samples and sequences of *Episoriculus* used for molecular analyses.

All the gene fragments were amplified with published primers [15,21–23]. PCR amplifications were carried out in a reaction volume mixture of 25 μl containing 12.5 μl 2×Taq Master Mix (Vazyme, Nanjing, China), 1 μl each primer, 1 μl genomic DNA, and 9.5 μl double-distilled water. PCR conditions for Cyt *b* amplifications consisted of an initial denaturing step at 94 °C for 5 min followed by 38 cycles of denaturation at 94 °C for 45 s, annealing at 49 °C for 45 s, extension at 72 °C for 90 s, and then a final extension step at 72 °C for 12 min. PCR conditions for the nuclear genes were basically the same as those of the Cyt *b*, but we changed the annealing temperature to 49-56 °C degrees (Apob: 49 °C; Brca1: 51 °C Rag2: 52 °C). We checked the PCR products on a 1.0% agarose gel and subsequently purified them using ethanol precipitation. Purified PCR products were directly sequenced using the BigDye Terminator Cycle Kit v 3.1 (Applied Biosystems, Foster City, CA, USA) and an ABI 310 Analyzer (Applied Biosystems).

To test the phylogenetic relationships within *Episoriculus*, we downloaded sequences of Cyt *b*, Apob, Brca1, and Rag2 for species of Nectogalini from GenBank for comparison (Table 2).

### Sequence analyses

All Cyt *b* sequences were aligned and examined, and screening for heterozygote nuclear gene fragments was performed in Mega 5 [24]. Referring to research articles on the same species [15,18–19], we used the three nuclear genes concatenated to analysis. Based on the amplified sequences in this study and downloaded sequences from NCBI, phylogenetic analyses were conducted on the following two datasets: 1) Cyt *b* genes; 2) three nuclear genes concatenated. We also conducted phylogenetic analyses based on each nuclear gene. Modeltest v 3.7 [24] was used to select the best fitting model of evolution, based on the Akaike Information Criterion in S1 Table. MrBayes v 3.1.2 [25] was used for the Bayesian analysis. Neogale vison was selected as the outgroup. Each run was carried out with four Monte Carlo Markov chains (MCMCs) and 10,000,000 generations for single gene datasets and 30,000,000 generations for concat-enated gene datasets. All runs were sampled every 10,000 generations. Convergences of runs were accepted when the average standard deviation of split frequencies fell be-low 0.01. Ultrafast bootstrap values (UFBoot) of ≽95 and posterior probabilities (PP) of ≽0.95 were considered strong support [26].

### Species tree and species delimitation

In this study, the *BEAST model was invoked in BEAST v1.7.5 using individual data analysis with complete genes in the Cyt *b*+nDNA dataset to construct species trees [27–28]. Model settings were selected with reference to the optimal replacement model of each gene (S1 Table). For each run, we selected the birth–death speciation model. The BEAST program was set up to run 30,000,000 generations, with statistical sampling every 5000 generations.

The Kimura-2-parameter (K2P) distances of the Cyt *b* gene between species were calculated in Mega 5 [24,29].

### Morphometric analyses

Specimens of *Episoriculus* used for morphological study were deposited in the Sichuan Academy of Forestry (SAF) and the Kunming Institute of Zoology (KIZ). Specimens were identified following Mittermeier and Wilson [3]. 56 complete skulls were used for geometric morphometric analyses, including 11 specimens of *E. caudatus*, 6 specimens of *E. sacratus*, 8 specimens of *E. umbrinus*, 17 specimens of *E. macrurus*, 7 specimens of *E. leucops*, and 3 specimens of *E. soluensis* (S2 Table).

According to the standard of Yang *et al*. [30], we measured the skulls of these specimens with a digital vernier caliper (accuracy 0.01 mm). Eleven measurements were taken: condylox incisive length (CIL), cranial height (CH), cranial breadth (CB), interorbital breadth (IBO), palatoincisive length (PIL), postpalatal length (PPL), maxillary breadth (MB), upper toothrow length (UTR), maximum width across the upper second molars (M_2_-M_2_), mandibular length (ML), and lower toothrow length (LTR). To evaluate the morphological variation between specimens, we performed a principal component analysis (PCA) in SPSS 19.0 (SPSS Inc., USA) using the log10-transformed craniodental measurements. Before analysis, the Kaiser–Meyer–Olkin (KMO) test and Bartlett’s sphere test were carried out. The KMO test was used to check the correlation and partial correlation between variables. Bartlett’s test determined if the correlation matrix was the unit matrix and the independent analysis method of each variable was invalid.

## Results and discussion

### Phylogenetic analysis

The Bayesian reconstruction using Cyt *b* (dataset 1) revealed eight monophyletic clades, corresponding to the following species: *Episoriculus macrurus, E. soluensis, E. leucops, E*. sp., *E. umbrinus, E. sacratus*, and *E. caudatus* (Fig. 2). *Pseudosoriculus fumidus* that once part of genus *Episoriculus* clustered with genus *Soriculus* with strong support (pp=0.98). *E. macrurus* clustered on an individual lineage and also splited from other species of genus *Episoriculus(pp=1.00). E. soluensis, E. leucops, E*. sp. clustered on an individual lineage respectively with strong support (pp=1.00). Both *E. umbrinus, E. caudatus*, and *E. sacratus* clustered on a lineage. Among them, *E. sacratus* was the first to differentiate with strong supported (pp=1.00). *E. umbrinus* and *E. caudatus* clustered on a common clade with significant support (pp=0.95)

**Fig 2.**
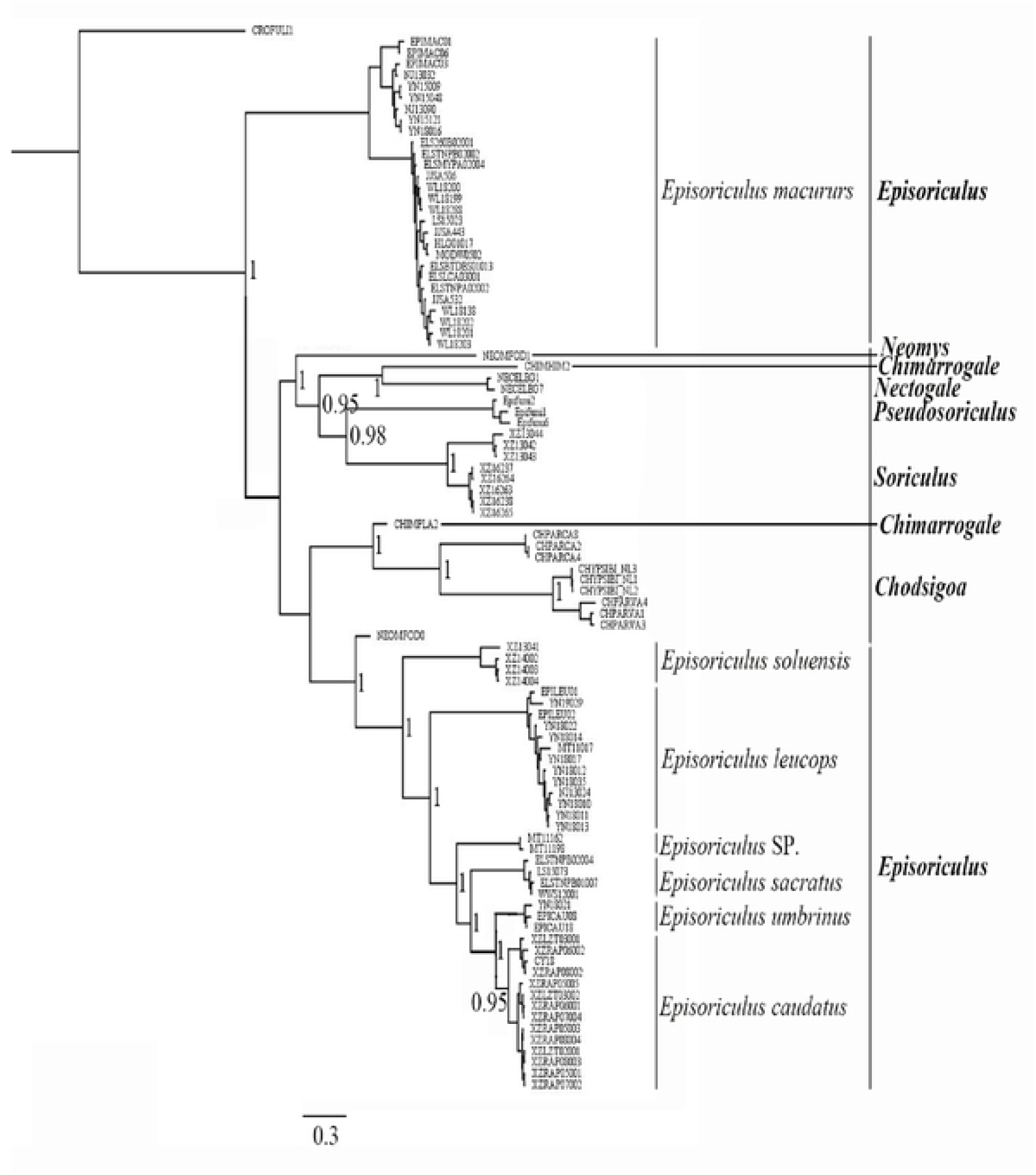
Bayesian phylogenetic analyses based on a dataset 1(Cyt *b*). Numbers at nodes. This is the Fig 2 legend.

The Bayesian reconstruction using the three nuclear genes also concatenated (dataset 2) revealed eight monophyletic clades (Fig. 3). In contrast to the Bayesian reconstruction using Cyt *b* (dataset 1), the topological structure was different. *Pseudosoriculus fumidus* clustered with genus *Chodsigoa* with strong support (pp=1.00). And all species of genus *Episoriculus* clustered on a common lineage*. E. macrurus* also clustered on an individual lineage with other species. *E. soluensis, E. leucops, E*. sp.*, E. umbrinus, E. sacratus*, and *E. caudatus* clustered on another lineage. Among them, *E. soluensis*, *E. leucops*, *E*. sp. also clustered on an individual lineage respectively with strong support (pp=1.00). Next, *E. sacratus* was the first to differentiate with strong supported (pp=1.00). *E. umbrinus* and *E. caudatus* also clustered on a common clade with significant support (pp=0.95).

**Fig 3.**
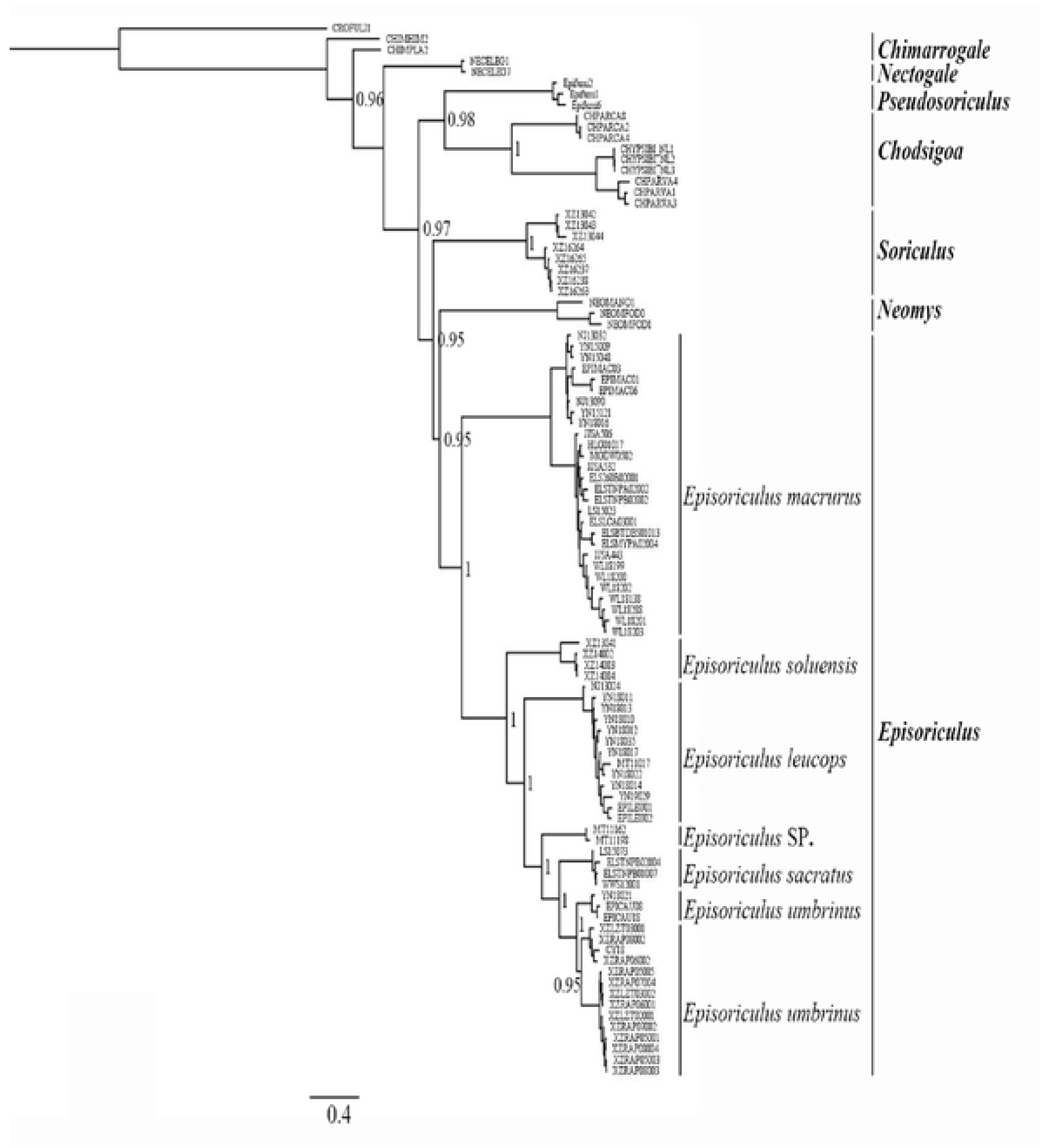
Bayesian phylogenetic analyses based on a dataset 2(three nuclear genes concatenated). Numbers at nodes. This is the Fig 3 legend.

### Species identification

The topology of *beast species trees differed slightly from those of the mitochondrial and nuclear gene trees (Fig. 4). In *beast species trees, the position of *Pseudosoriculus fumidus* was the same as in the mitochondrial tree, however, is not well support(pp=0.87). *E. soluensis, E. leucops*, and *E. macrurus* clustered on an individual lineage with strong support respectively. *E. umbrinus* and *E. caudatus* also *c*lustered on a common clade with significant support(pp=0.95). But *E. sacratus* and *Episoriculus* sp. got strong supported.

**Fig 4.**
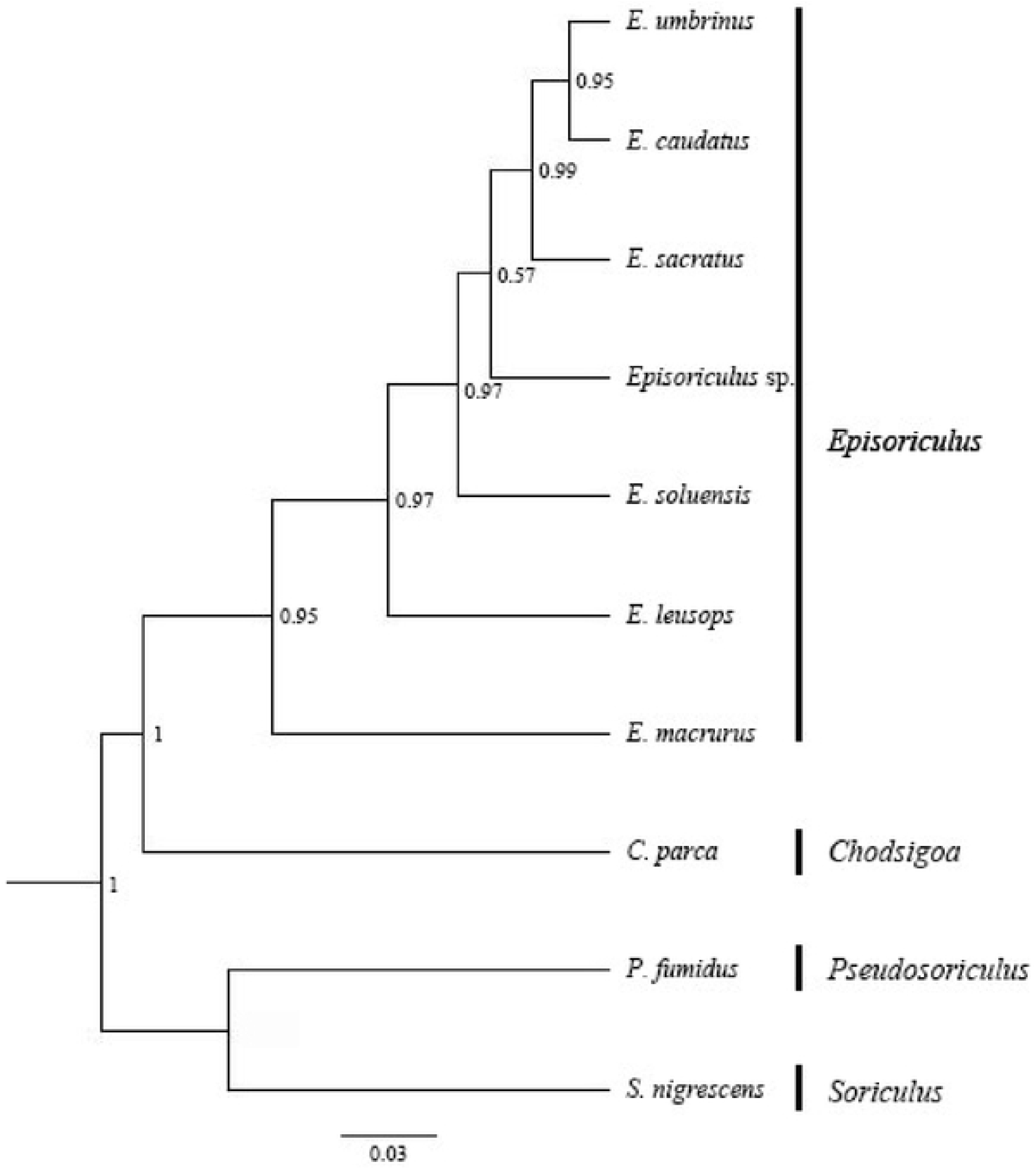
Results of species delimitation nDNA gene species trees using the BEAST model. Node numbers indicate Bayesian posterior probabilities supporting. This is the Fig 4 legend.

The Kimura-2-parameter distances between the species of *Episoriculus* ranged from 0.021 to 0.189 (Table 3). The average Kimura-2-parameter distance between *Pseudosoriculus fumidus* and other species of the genus *Episoriculus* was 0.165. The average Kimura-2- parameter distance between *E. macrurus* and other species of the genus *Episoriculus* was 0.136. The Kimura-2-parameter distance between *E. caudatus* and *E. sacratus* was 0.033. The Kimura-2-parameter distance between *E. caudatus* and *E. umbrinus* was 0.021. The Kimura-2- parameter distance between *E. sacratus* and *E. umbrinus* was 0.028.

**Table 3.** The Kimura-2-parameter distances between *Episoriculus* species based on the Cyt *b* gene.

| No. | Species | 1 | 2 | 3 | 4 | 5 | 6 |
| --- | --- | --- | --- | --- | --- | --- | --- |
| 1 | *E. macrurus* |  |  |  |  |  |  |
| 2 | *E. sacratus* | 0.125 |  |  |  |  |  |
| 3 | *E. umbrinus* | 0.133 | 0.028 |  |  |  |  |
| 4 | *E. caudatus* | 0.134 | 0.033 | 0.018 |  |  |  |
| 5 | *E. leucops* | 0.146 | 0.110 | 0.111 | 0.124 |  |  |
| 6 | *E. soluensis* | 0.132 | 0.077 | 0.079 | 0.103 | 0.119 |  |
| 7 | *E. fumidus* | 0.144 | 0.163 | 0.169 | 0.178 | 0.189 | 0.148 |

### Morphological analyses

Morphological index measurement data are shown in Table 5–6, complete measurement data are provided in S2 Table. Body length (HBL), tail length (TL), hind foot length (HFL), ear height (EL), and body weight (BW) of all 56 specimens with intact skulls were counted (Table 4). The tail length of *Episoriculus macrurus* was 71–106mm, while those of the congeners ranged from 46 to 82.5mm. The HFL of *E. macrurus* was usually 15mm, while other species usually 12–13mm. *E. macrurus* had minimum HBL/TL ratio of 0.64, and its TL was close to 1.5 times of its HBL, while the HBL of other congeners was approximately equal or greater than the TL. Meanwhile, *E. leucops* had the largest body, and *E. sacratus* differed from *E. caudatus* and *E. sacratus* by HBL/TL.

**Table 5.** The results of body morphologic measurements.

| Species | N | BW | HBL | TL | HFL | HBL / TL |
| --- | --- | --- | --- | --- | --- | --- |
| *E. caudatus* | 14 | 5.000±0.327 | 56.750±1.858 | 54.750±0.491 | 12.125±0.125 | 1.04 |
| *E. sacratu*s | 6 | 5.666±0.211 | 57.134±0.632 | 62.166±2.249 | 13.500±0.671 | 0.92 |
| *E. umbirnu*s | 9 | 5.000±0.239 | 61.333±0.816 | 60.222±0.547 | 13.000±0.231 | 1.01 |
| *E. leucops* | 7 | 7.714±0.286 | 62.714±0.056 | 60.143±1.534 | 13.143±0.143 | 1.04 |
| *E. macrurus* | 17 | 5.000±0.413 | 58.158±1.732 | 91.333±1.201 | 15.667±0.333 | 0.64 |
| *E. soluensis* | 4 | 6.000±0.210 | 64.333±0.666 | 61.333±0.088 | 12.000±0.531 | 1.04 |

**Table 6.** External and craniomandibular measurements including the mean values and ranges of six.

| Species | N | CIL | IOB | CB | BH | MB | PIL | PPL | UTR | M2-M2 | ML | LTR |
| --- | --- | --- | --- | --- | --- | --- | --- | --- | --- | --- | --- | --- |
|  |  | Mean±S.D |  |  |  |  |  |  |  |  |  |  |
| *E. caudatus* | 14 | 18.327 | 3.311 | 8.769 | 5.907 | 1.391 | 8.104 | 8.067 | 8.1464 | 4.525 | 11.258 | 7.2657 |
|  |  | ±0.057 | ±0.044 | ±0.374 | ±0.068 | ±0.018 | ±0.059 | ±0.039 | ±0.058 | ±0.038 | ±0.056 | ±0.058 |
| *E. sacratu*s | 6 | 18.206 | 3.430 | 8.578 | 6.030 | 1.381 | 7.840 | 8.035 | 7.773 | 4.565 | 11.036 | 6.981 |
|  |  | ±0.0593 | ±0.052 | ±0.082 | ±0.100 | ±0.237 | ±0.056 | ±0.103 | ±0.073 | ±0.029 | ±0.090 | ±0.046 |
| *E. umbirnu*s | 8 | 18.286 | 3.523 | 8.513 | 5.628 | 1.371 | 8.208 | 8.126 | 8.052 | 4.457 | 11.1789 | 7.3933 |
|  |  | ±0.070 | ±0.054 | ±0.045 | ±0.045 | ±0.024 | ±0.063 | ±0.088 | ±0.046 | ±0.049 | ±0.077 | ±0.0763 |
| *E. leucops* | 7 | 19.307 | 3.708 | 9.068 | 5.858 | 1.391 | 8.658 | 8.876 | 8.416 | 4.564 | 11.693 | 7.354 |
|  |  | ±0.111 | ±0.377 | ±0.448 | ±0.046 | ±0.341 | ±0.036 | ±0.797 | ±0.082 | ±0.062 | ±0.130 | ±0.147 |
| *E. macrurus* | 18 | 17.577 | 3.309 | 8.456 | 5.761 | 1.290 | 7.633 | 7.813 | 7.565 | 4.184 | 10.618 | 6.781 |
|  |  | ±0.060 | ±0.191 | ±0.057 | ±0.050 | ±0.012 | ±0.051 | ±0.079 | ±0.056 | ±0.045 | ±0.051 | ±0.036 |
| *E. soluensis* | 4 | 17.896 | 3.603 | 9.327 | 5.763 | 1.706 | 8.067 | 7.976 | 7.767 | 4.873 | 12.320 | 7.497 |
|  |  | ±0.003 | ±0.052 | ±0.097 | ±0.233 | ±0.328 | ±0.296 | ±0.024 | ±0.084 | ±0.003 | ±0.055 | ±0.084 |

### species including one cryptic species of *Episoriculus*

Bartlett’s test rejected the null hypothesis (χ 2 = 568.01, P = 0.000), which indicated that the measurements were consistent with factor analysis. KMO was 0.823, indicating that there was a strong correlation among the variable data of skull morphology, which was suitable for factor analysis. Two principal components, explaining 74.69% of the morphological variation, were extracted from the analysis. Its factor loading values were all positive and most were greater than 0.95 (Table 7), indicating that it was mainly related to the overall size of the skull. The first principal component (PC1) had the highest degree of explanation (61.022%), and the features with factor load > 0.9 included PIL, ML, CIL, UTR, and LTR.

**Table 7.**
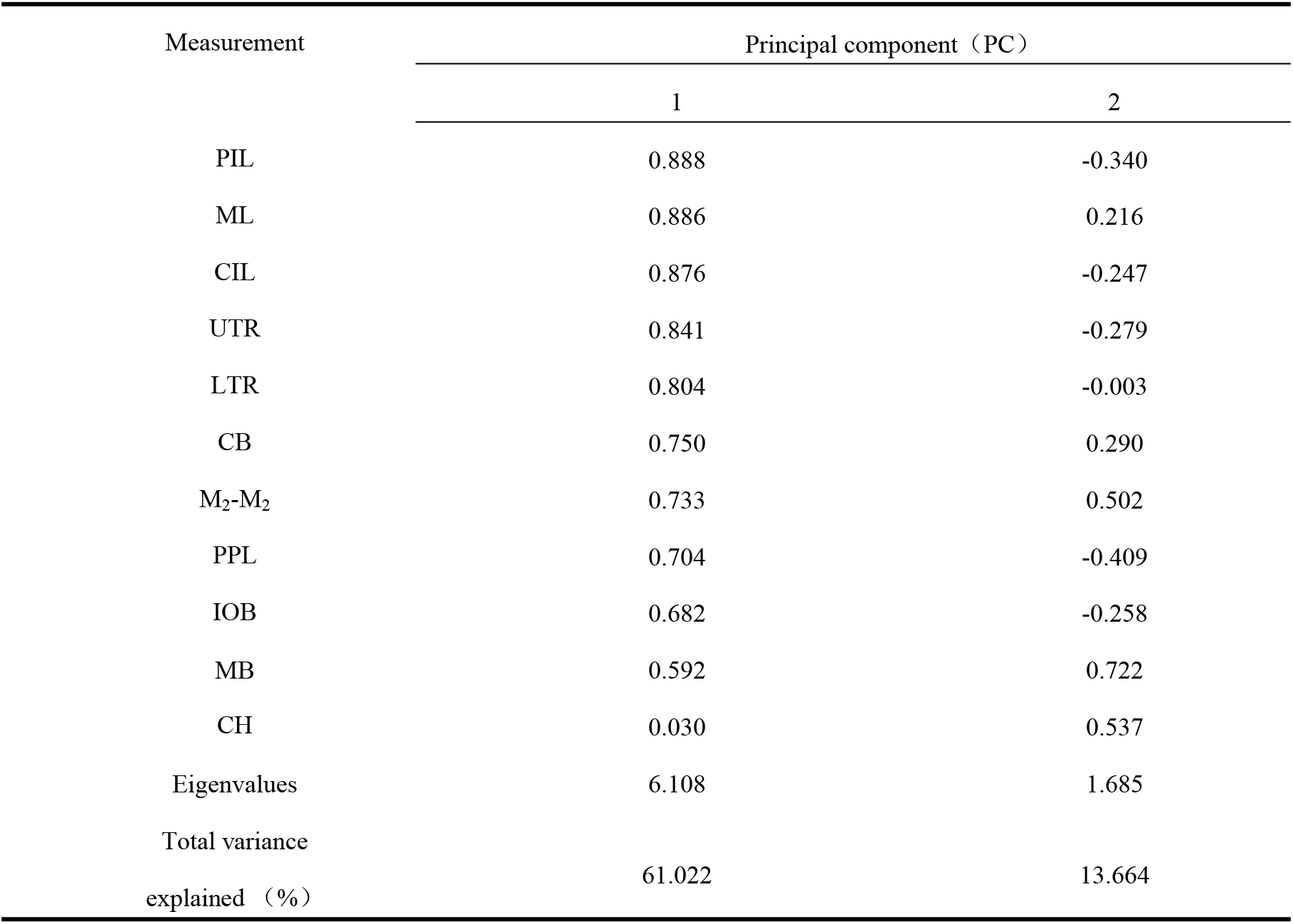
Character loading, eigenvalues, and percent variance explained on the two components of PCA of cranial measurements of *Episoriculus*.

| Measurement | Principal component (PC) |  |
| --- | --- | --- |
|  | 1 | 2 |
| PIL | 0.888 | -0.340 |
| ML | 0.886 | 0.216 |
| CIL | 0.876 | -0.247 |
| UTR | 0.841 | -0.279 |
| LTR | 0.804 | -0.003 |
| CB | 0.750 | 0.290 |
| M<sub>2</sub>-M<sub>2</sub> | 0.733 | 0.502 |
| PPL | 0.704 | -0.409 |
| IOB | 0.682 | -0.258 |
| MB | 0.592 | 0.722 |
| CH | 0.030 | 0.537 |
| Eigenvalues | 6.108 | 1.685 |
| Total variance explained (%) | 61.022 | 13.664 |

Using PCA1 and PCA2 maps (Fig. 5), *E. leucops, E. macrurus*, and *E. soluensis* in concentrated regions, indicating substantial morphological differences. *E. macrurus* was plotted in the negative region of PC2, while those of the other species were mainly in the positive region. *E. leucops* was plotted in the negative region of PC1 and the positive region of PC2. While *E. soluensis* was plotted in on the positive regions of the PC1 and PC2. However, *E. sacratus*, *E. umbrinus*, and *E. caudatus* were not distinguished well and mixed together. Using LAD maps (Fig 6), *E. sacratus*, *E. umbrinus*, and *E. caudatus* were distinguished well. Among them, two of nine individuals of *E. caudatus* were wrongly judged as *E. sacratus*, and one individual of *E umbrinus* was wrongly identified as *E. sacratus*.

**Fig 5.**
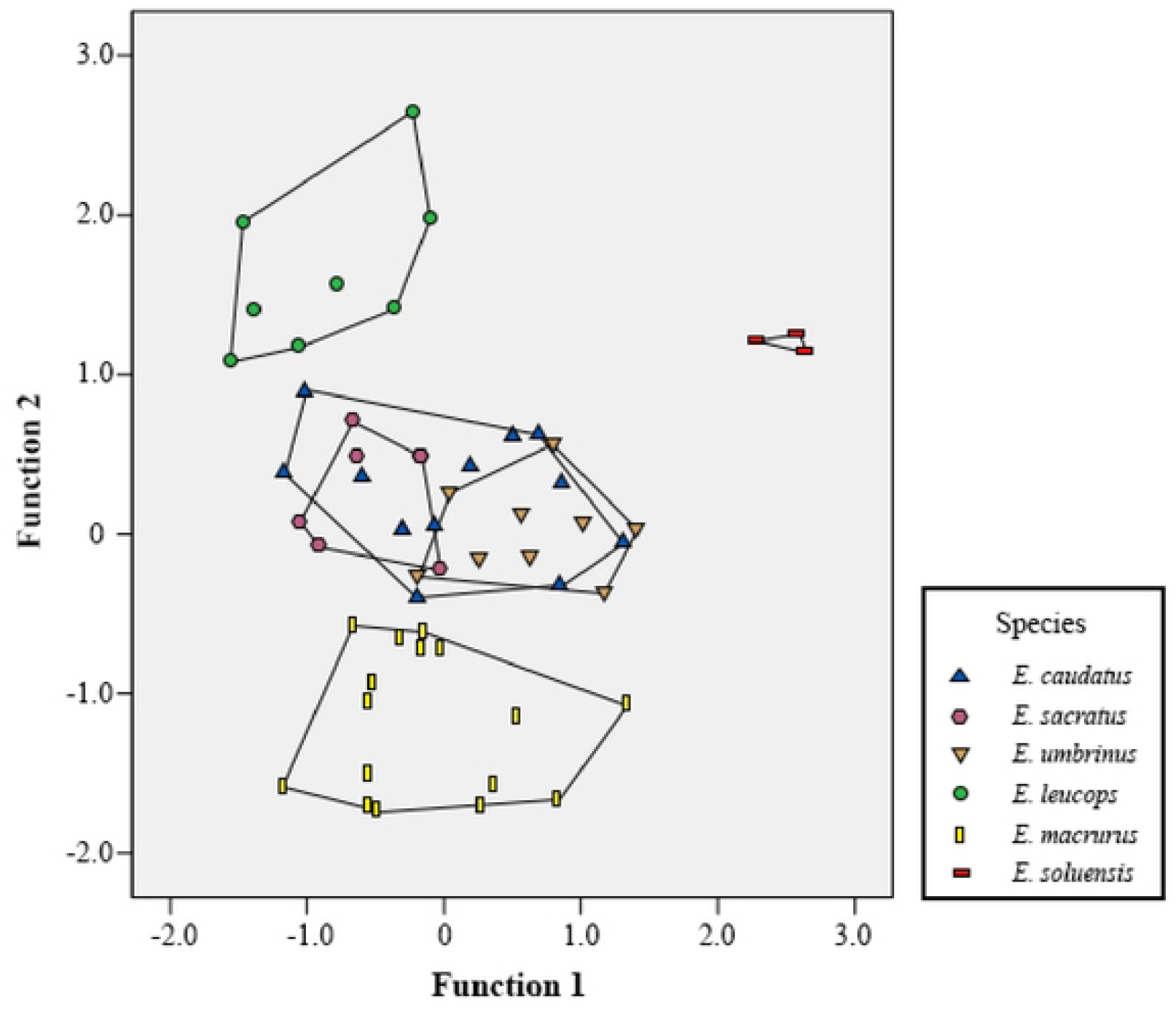
Results of principal component analysis of *Episoriculus* based on the 19 log10- transformed craniodental measurements. This is the Fig 5 legend.

**Fig 6.**
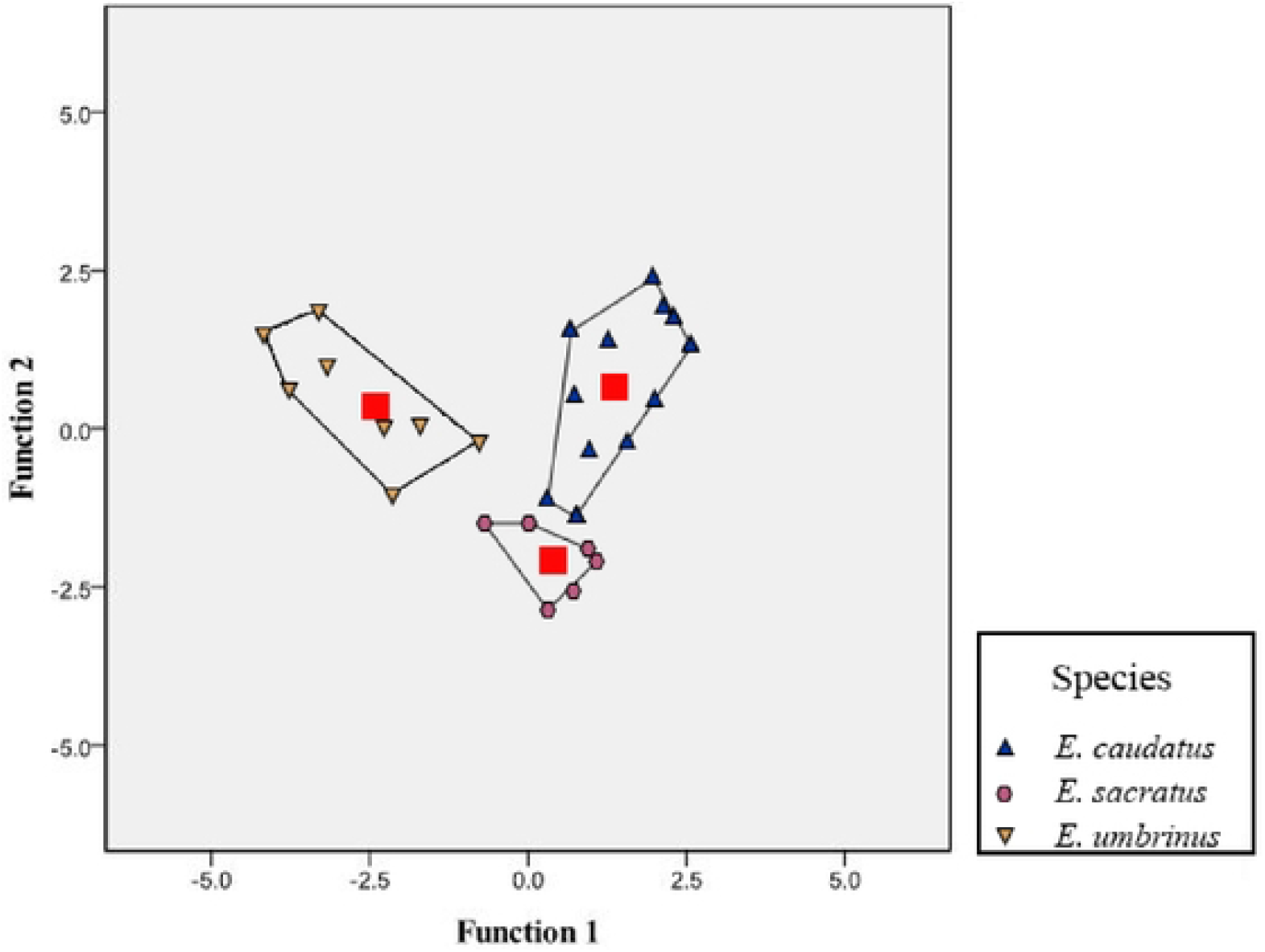
Results of principal component analysis of *Episoriculus* based on the linear discriminant analysis. This is the Fig 6 legend.

The skull characteristic of some species of *Episoriculus* (Fig. 7) showed that the braincase of *E. macrurus* was very dome-shaped, it had the smallest width and orbital distances, and rostrum. Meanwhile, the upper unicuspids of *E. macrurus* were quadrangle and wider than longer, while the upper unicuspids of other species were usually rectangular.

**Fig 7.**
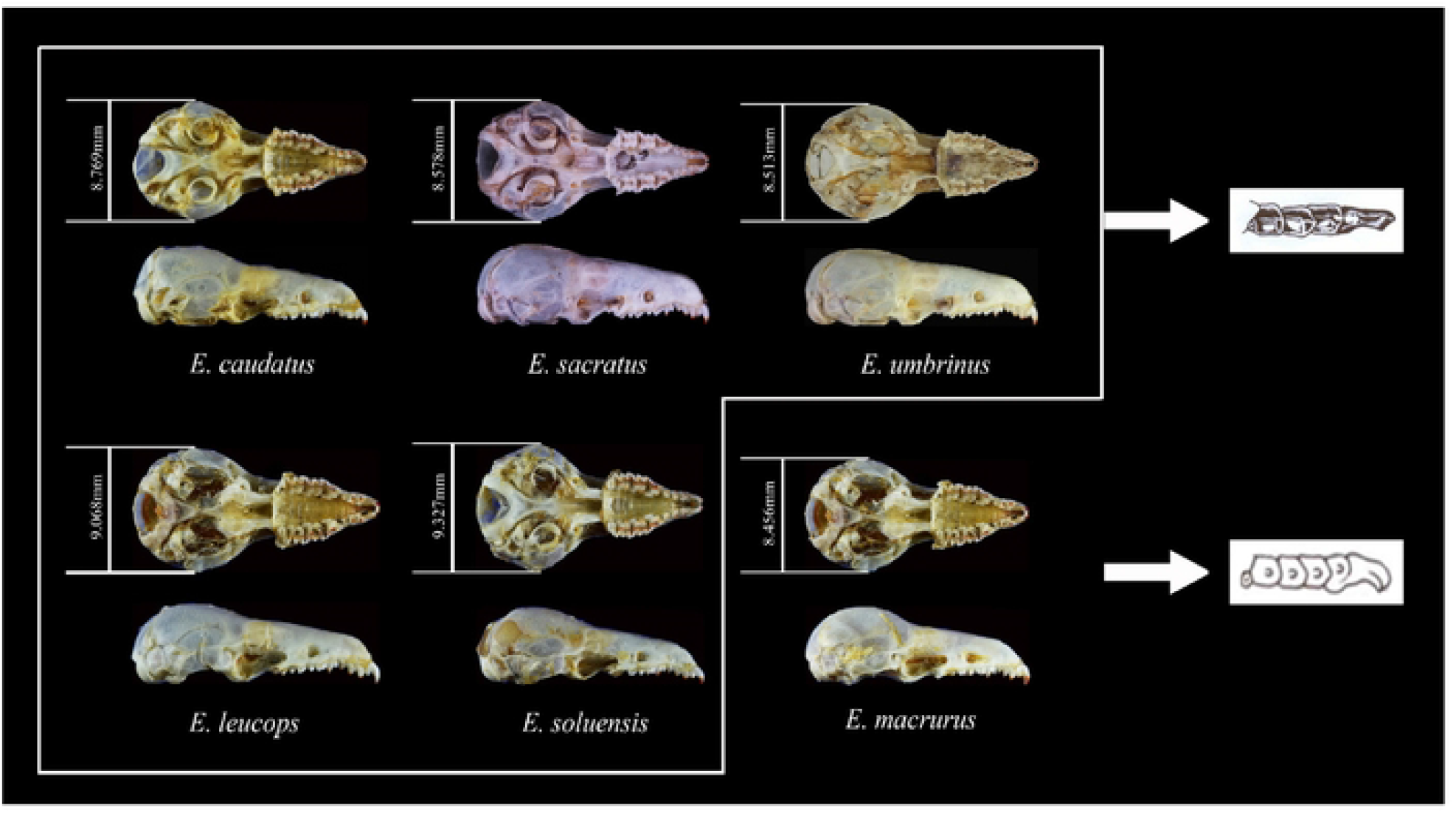
The comparison of skulls in genus *Episoriculus*. This is the Fig 7 legend.

## Discussion

### Differences between mitochondrial and nuclear genes

Conflicting phylogenetic signals between mitochondrial and nuclear genes have been reported in many studies [31]. There are many reasons for this conflict, such as differences in genetic background [32], Ancient hybridization [33], incomplete lineage sorting [34], adaptive evolution, and burst Formula speciation [35]. In this study, we also found that the mitochondrial gene tree is inconsistent with the nuclear gene tree.

In this study, we considered that mitochondrial genes can provide a lot of valuable information in solving the phylogenetic relationship of *Episoriculus*. However, neither mtDNA nor nDNA alone can completely solve the phylogenetic problem of *Episoriculus*, and the use of data from these two different genetic pathways has achieved good results; species tree construction in the anadromous framework also resulted in a consistent topology with high statistical approval ratings. Therefore, it is necessary to combine mitochondrial gene and nuclear gene information when solving the phylogenetic problem of *Episoriculus*

### Species of *Episoriculus*

For a long time, many scholars regarded *E. sacratus* and *E. umbrinu* as subspecies of *E. caudatus* [1,7,17]. Motowaka *et al*. [14] promoted the recognition of both *E. caudatus* and *E. sacratus* based on differences in karyotype, and considered *E. umbrinus* as a subspecies of *E. sacratus*. Nevertheless, Mittermeier and Wilson [3] elevated *E. umbrinus* to a species without any verification. In the study, phylogenetic analysis and species identification supported that *E. umbrinus*, *E. caudatus* and *E. sacratus* are independent species, and that the three species also have morphological differences.We recommend a new cryptic species collected in low elevation areas in Motuo County, Tibet (*Episoriculus* sp.). However, we only collected two samples and could not accurately describe their morphology.

Next, Abramov *et al*. [2] promoted *E. soluensis* as an independent species, which was a synonym of *E. caudatus*. And the type locality of *E. soluensis* is Pokhara, Nepal in the southern Himalayas (Abramov). Based on comparison of phylogenetic analysis, we confirmed that four specimens collected from Yadong and Nyalam is *E. soluensis*, and it constitutes a new record of the species occurrence in China.

Motokawa and Lin [13] considered that *E. baileyi* is a valid species of genus *Episoriculus*, and it is a large species with a tail shorter than or as long as the head and body length, and is distinguished from other *Episoriculus* species in having the combination of robust first upper incisor, long rostrum and upper unicuspids row, large tympanic ring, and high ascending ramus of mandible. Therefore, we agreed the opinion of Motokawa and Lin (2005). However, its taxonomic status needs to be further determined in phylogenetic study.

Hoffmann [7], Motokawa and Lin [13] pointed out that *P. fumidus* has a unique morphology. Abramov *et al*. [2] erected the new genus *Pseudosoriculus* for the species. Our analysis also supported new classification status of *Pseudosoriculus*, and confirmed that *P. fumidus* does not belong to genus *Episoriculus*.

In the study, our phylogenies analysis consistently rooted *E. macrurus* as an individual lineage*. E. macrurus* has a large genetic distance from other species of genus *Episoriculus*, Morphologically*, E. macrurus* has the longest tail, and also differences substantially from its congeners in its skull and teeth between *E. macrurus* and other species of genus *Episoriculus*. In conclusion, we erected a new subgenus *Longacauda* including *E. macrurus*, while *E. caudatus*, *E. sacratus*, *E. umbrinus*, *E. soluensis*, *E. leucops*, *E. baileyi*, and *Episoriculus* sp. formed another subgenus: *Episoriculus*.

**Table 8.**
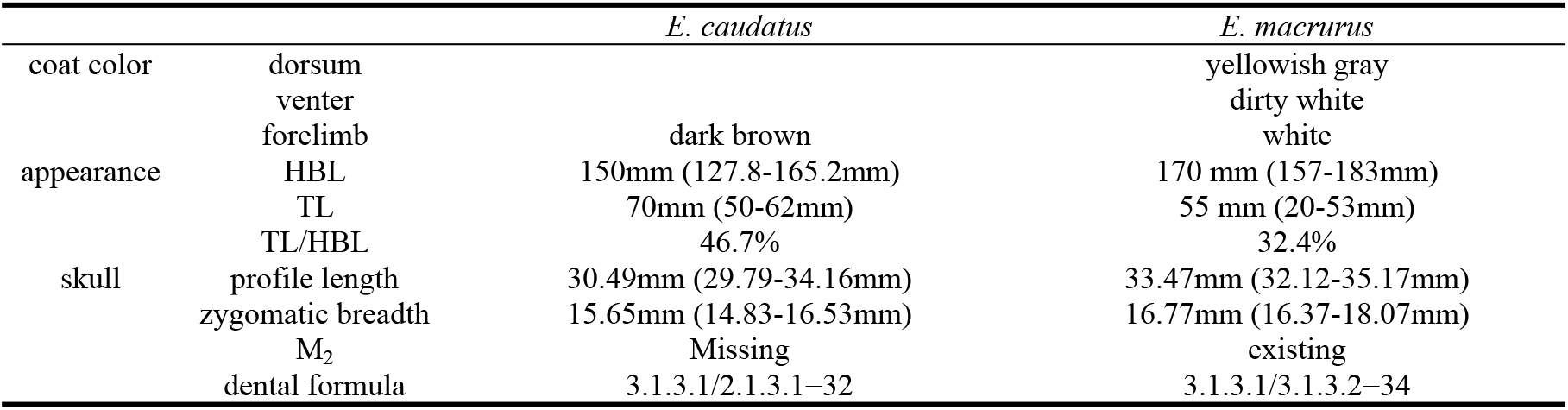
Significant morphological differences of *E. caudatus* and *E. macrurus*.

### Systematic account

Family: Soricidea

Subfamily: Soricinae

Genus: *Episoriculus*

Subgenus: *Longacauda* Liu, Jin, Wang, Liao, Yue, and Liu, **subgen. nov.**

Type species: *Episoriculus macrurus* (as published in The Fauna of British India including Ceylon and Burma. Blanford, 1888).

*Etymology—*The subgeneric name, negative compound word, combining the word *Longa* meaning long and the word *cauda* meaning tail in Latin.

*Diagnosis —*Long tail and long hindfoot. Tail length is 71–106mm and close to 1.5 times of body length. The hindfoot is about 15mm. Braincase is very dome-shaped, rostrum is small, and the upper unicuspid quadrangular, and wider than longer.

This subgenus includes one species: *E. macrurus*.

## Conclusions

Based on molecular and morphological analyses, the genus *Episoriculus* contains two subgenus: *Longacauda* and *Episoriculus*, and including eight species: *E. baileyi*, *E. caudatus*, *E. leucops, E. macrurus, E. sacratus, E. soluensis, E. umbrinus, and Episoriculus* sp.

## Acknowledgments

This research was funded by the National Natural Science Foundation of China (31970399). We are grateful to Robert Murphy to answer and correct scientific questions. We are grateful to Xuming Wang, Yingting Pu, and Jiao Qing for their assistance with this study.

## Supporting information

S1 Table. Primers used in PCR and sequencing.

S2 Table. External and selected cranial measurements of *Episoriculus* species.

